# DyNeuMo Mk-1: Design and Pilot Validation of an Investigational Motion-Adaptive Neurostimulator with Integrated Chronotherapy

**DOI:** 10.1101/2020.09.10.292284

**Authors:** Mayela Zamora, Robert Toth, Francesca Morgante, Jon Ottaway, Tom Gillbe, Sean Martin, Guy Lamb, Tara Noone, Moaad Benjaber, Zachary Nairac, Devang Sehgal, Timothy G. Constandinou, Jeffrey Herron, Tipu Z. Aziz, Ivor Gillbe, Alexander L. Green, Erlick A. C. Pereira, Timothy Denison

## Abstract

There is growing interest in using adaptive neuromodulation to provide a more personalized therapy experience that might improve patient outcomes. Current implant technology, however, can be limited in its adaptive algorithm capability. To enable exploration of adaptive algorithms with chronic implants, we designed and validated the ‘Picostim DyNeuMo Mk-1’ (DyNeuMo Mk-1 for short), a fully-implantable, adaptive research stimulator that titrates stimulation based on circadian rhythms (e.g. sleep, wake) and the patient’s movement state (e.g. posture, activity, shock, free-fall). The design leverages off-the-shelf consumer technology that provides inertial sensing with low-power, high reliability, and relatively modest cost. The DyNeuMo Mk-1 system was designed, manufactured and verified using ISO 13485 design controls, including ISO 14971 risk management techniques to ensure patient safety, while enabling novel algorithms. The system was validated for an intended use case in movement disorders under an emergency-device authorization from the Medicines and Healthcare Products Regulatory Agency (MHRA). The algorithm configurability and expanded stimulation parameter space allows for a number of applications to be explored in both central and peripheral applications. Intended applications include adaptive stimulation for movement disorders, synchronizing stimulation with circadian patterns, and reacting to transient inertial events such as posture changes, general activity, and walking. With appropriate design controls in place, first-in-human research trials are now being prepared to explore the utility of automated motion-adaptive algorithms.

## 1 Introduction

As the field of adaptive neuromodulation is rapidly evolving, a key question is what signals to use for adapting stimulation delivery; arguably the current emphasis is on using bioelectric signals to inform the control algorithm [1–4]. As the leading commercial system, the Neuropace RNS is approved in the U.S. for refractory epilepsy [5]. While promising, the ultimate benefit of the responsive stimulation for epilepsy is still debated, and refinement of the algorithmic approach remains an active area of study [6]. Likewise, in the field of movement disorders, particularly Parkinson’s disease, adaptive stimulation has shown promise for improving outcomes while lowering energy use [3, 4]. However, the signals recorded from sub-cortical targets are 1) relatively small (1 µVrms) [3, 4, 7], 2) prone to artefacts from stimulation, cardiac signals and motion [8, 9], and 3) the optimal configuration of algorithms are still debated and might prove complex for programming [10]. In addition, the resolution of small bioelectric signals in the presence of stimulation puts significant constraints on the relationship between sensing and stimulation electrodes, which can severely limit the therapy options [3, 4, 11, 12]; recent work to bypass these constraints potentially compromise the safety of the tissue-electrode interface due to leakage currents and single-fault errors [13].

There are several alternatives to bioelectrical signals which might be used to adjust a stimulator. For example, adapting stimulation with time could be a simple, yet impactful feedforward approach to therapy optimization. As pharmaceuticals have been shown to exhibit sensitivity to timing [14], implantable devices might also benefit from exploiting rhythmicity linked to disease processes [15, 16]. Specifically, time-varying disease processes might be synchronized with stimulation adjustments, thereby implementing chronotherapy through use of the embedded real-time clock in bioelectronic circuits [17]. As another algorithm input source, inertial sensors can also be used to obtain an estimate of the patient or symptom state as a feedforward method to adjust stimulation. The widespread adoption of inertial sensing in consumer wearable electronics has resulted in many features ideal for use in implantable closed-loop neuromodulation systems: 1) low power (order of 10 µW), 2) high reliability and shock immunity, and 3) embedded “digital motion classifiers” that facilitate state estimation [18]. Inertial sensing has already been applied in medical implants to automatically titrate stimulation parameters. Notable examples include activity-based tuning of cardiac pacemakers [19], and posture responsive adjustment of stimulation for spinal cord stimulation for chronic pain [20]. Investigational work using the Activa PC+S also demonstrated the potential utility of inertial sensing for deep brain stimulation (DBS) applications such as essential tremor [21] and Parkinson’s disease [22]. Despite the potential research and therapeutic opportunities enabled by integrating inertial and circadian adaptive functionality into neuromodulation systems, there are no such devices currently available for in-human research.

In this paper, we introduce the Dynamic Neuro Modulator Mark 1 (DyNeuMo Mk-1), a cranially-mounted circadian- and motion-adaptive neurostimulator for use in first-in-human investigational studies exploring circadian- and inertial-sensing based closed-loop therapies. The system is based on the predicate Picostim system manufactured by Bioinduction [23], that provides several advantages as a therapy research platform. The small device size of 7 cc and recharge capability also allows for flexible use throughout the body. Unlike existing deep brain stimulation devices, which are implanted in the chest cavity with electrode leads routed through the neck, the Picostim systems use a cranial mounted design. The surgical procedure for device placement has some similarities to cochlear implant devices, and the infection risk might potentially be lowered compared to existing DBS procedures [24]. Most notably, the cranial-mounting avoids tunnelling leads through the neck, that could reduce the risk of lead wire breakage or fibrosis in the surrounding tissue, a potential cause of stiffness and pain [25]. The Picostim firmware and software can also be modified to enable novel adaptive algorithms, including time- and inertial-based inputs, which supports its utility as a flexible therapy research tool [26, 27].

The DyNeuMo Mk-1 added research subsystems to the predicate Picostim design. To support first-in-human research, we used ISO 13485–compliant design controls throughout the project. The paper will follow a similar structure to a typical medical device design flow, starting with the assessment of our device requirements motivated by anticipated user needs and risk management. We will then discuss in detail the implementation of our design before demonstrating the system’s functionality through both verification testing and a subacute test of adaptive algorithms in a subject with cervical dystonia. Future research projects for system validation are briefly outlined, as well as a discussion of the advantages and limitations of the implemented approach. The circadian- and inertial-focused research stimulator will expand the possible research space for human feasibility studies, providing an alternative method for adaptive, patient-specific therapies.

## 2 Design Requirements and Implementation

We designed the DyNeuMo Mk-1 to be used as a research system for exploring how we might improve therapies with automated algorithms. The system-level requirements are summarized in Table 1. From an architecture perspective, the DyNeuMo Mk-1 was implemented using the physiologic control model of Fig. 1 [1]. To summarize, our aim is to supplement the selection of stimulation parameters using manual and timer-scheduled adjustments with the addition of motion-adaptive changes. This can be considered an additional response loop that adjusts stimulation based on characteristic motion profiles. Using this framework, we present the key attributes of the design, and how the user engages the adaptive stimulation functionality. The implementation of the system block diagram and its decomposition into sub-components is shown in Fig. 2, which illustrates the control flow, signal routing, and hardware embodiment of the DyNeuMo Mk-1.

**Table 1.** System-level specifications for the DyNeuMo-Mk1 investigational research system

| **User Needs** |  |
| --- | --- |
| Predicate Therapy Support | The research system must support existing stimulation parameters for therapy delivery (amplitude, frequency, pulse width) |
| Supported Therapy Research | Deep brain stimulation, chronic pain (spinal cord), incontinence and bladder control (sacral, pudendal nerve), gastroparesis (enteric) |
| Adaptive Sensing Scheme | Circadian scheduler – temporal program selection in 30-minute epochs, repeated on a 24-cycle. The program can be activated for a sub-section of each epoch<br>Inertial accelerometer (three axis) – with DC accuracy for posture detection and AC capability for activity, tremor, gait, shocks and free-fall – flexibility for configuration to specific therapy needs |
| Algorithm Training Support | Ability to stream data for classifier training |
| Algorithm Power Allowance | The adaptive algorithm must draw no more than 20% of the nominal therapy power (e.g. 80 μW for deep brain stimulation) |
| Algorithm Latency | < 20 ms from event detection to stimulation adjustment |
| Algorithm Verification | Minimal-risk verification procedure for the algorithm |
| **Risk Mitigations** |  |
| Stimulation (Actuation) Limits | Pre-defined limits on the stimulation level to ensure patient safety; this includes transition ramps between stimulation program settings for tolerance (e.g. avoid paresthesia) |
| State Monitoring and Alerts | Algorithm state clearly shown on patient interface, ability to enable/disable with a button press; data logger for algorithm transitions for issue resolution |
| Fallback Mode | Pre-defined stimulation program for emergency exit from algorithm; disengagement of the automated algorithm during recharge |
| Physiological Dynamics | Stimulation timing interlocks to avoid inadvertent rapid transitions at classification boundaries |

**Fig. 1.**
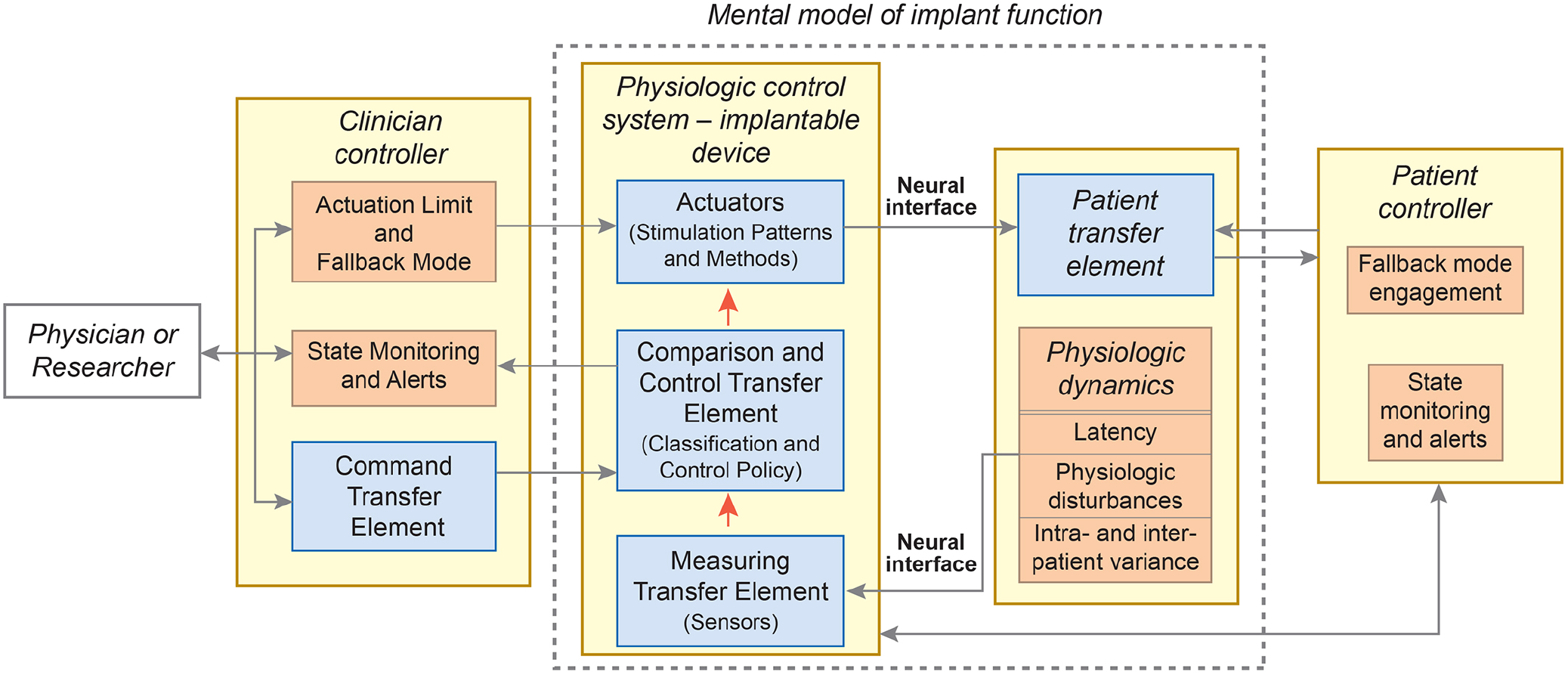
System block-diagram using the IEC 60601-1-10 physiologic control framework [1]. Blue boxes are derived from user needs, while tan boxes are derived from risk mitigations. Both sources of design inputs inform the system specifications for the DyNeuMo Mk-1.

**Fig. 2.**
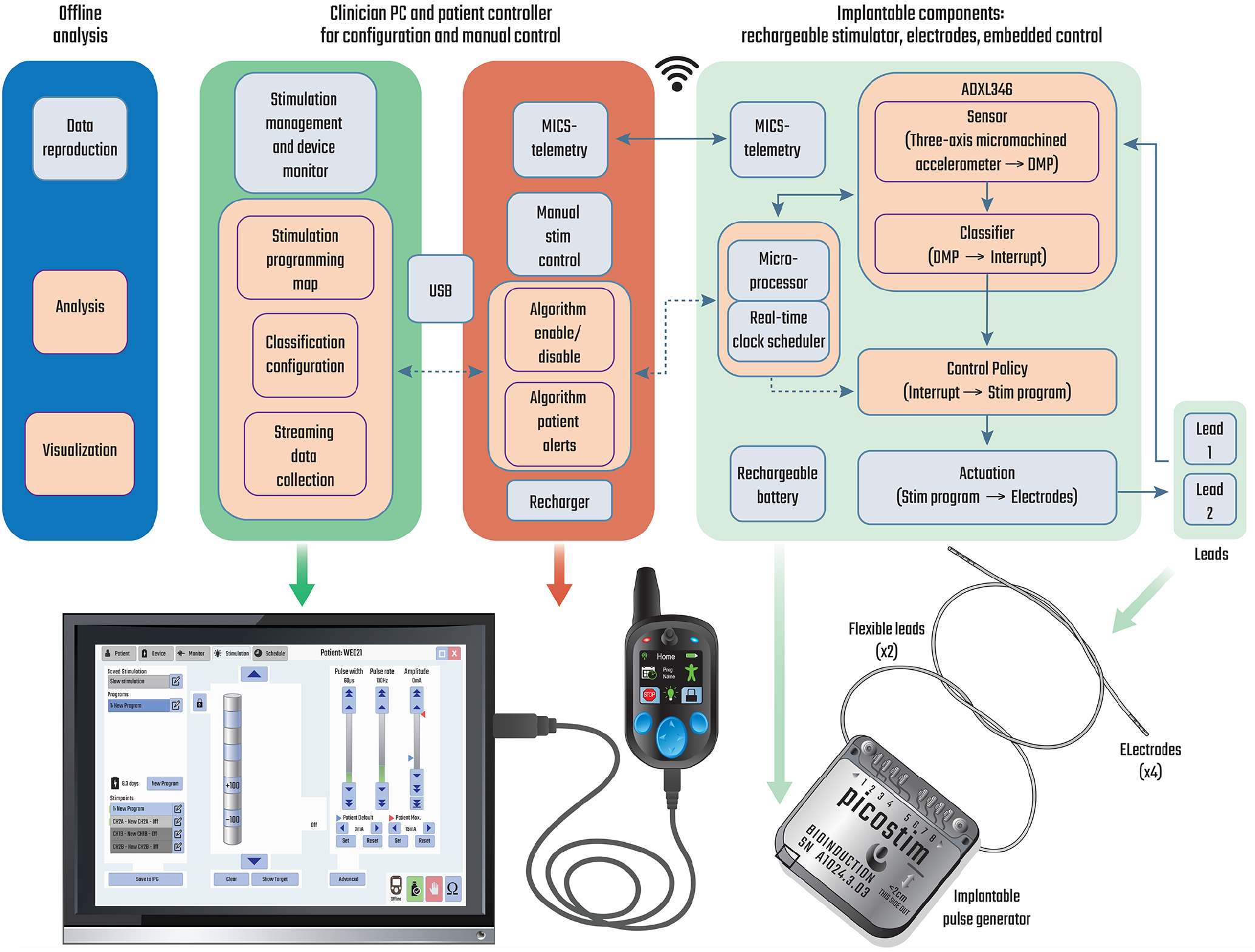
**Top:** system block diagram of the DyNeuMo Mk-1, illustrating the baseline functionality provided by the predicate Picostim system (light blue) and the algorithmic additions (tan). **Bottom:** realisation of the research platform. Hardware elements of the toolchain are largely reused from the predicate system to leverage their existing verification as part of the device quality management system. The implantable pulse generator can be configured and monitored via MICS–band telemetry from the clinician programmer. In-clinic programming of the handheld patient controller takes place over a USB link to the clinician tablet. Research subjects use the handheld controller for monitoring charge level, and manual adjustments to the stimulation program.

### 2.1 User Needs: The Mental Model for Operation and Preservation of Existing Actuation

As a first design requirement, our research tool must preserve the stimulation capabilities of predicate therapy systems, to ensure there is no compromise to clinical care options. This approach is consistent with other state-of-the-art research tools provided for therapy research [28, 29]. The DyNeuMo Mk-1 provides stimulation capability equivalent to predicate deep brain, chronic pain, sacral nerve (incontinence), and gastric stimulators, based on publicly-available manufacturer specifications.

As a general research tool, we aim to support a variety of potential use-cases. Motion-based states of interest include tremor (oscillations), general activity, gait and freezing, absolute posture, falls, and transient shocks. The detection of these motion states can be applied by researchers exploring improved therapies for postural and gait instability in Parkinson’s disease [30], transient stress events in mixed incontinence [31], posture effects such as orthostatic hypertension [32], and titration of stimulation through circadian (sleep-wake) cycles [33]. In addition to automated stimulation titration, inertial sensing also provides diagnostic information on patient activity without burdening the user with added instrumentation. Finally, the sensor also provides an alternative input method for the patient to discretely interact with their device through explicit motor inputs, such as tap-activation.

The practical implementation of a motion-adaptive stimulator motivates additional design requirements. To help train and program the classifier, we need a method to gather individual patient data and configure the algorithm based on their specific characteristics. In addition, a control policy is required to map the outcome of motion classification to the desired stimulation state. To minimize the impact on device longevity or avoid increasing recharge burden, the addition of the algorithm must not significantly increase the power consumption of the system compared to baseline therapy, e.g. roughly < 400 µW for bilateral stimulation in the treatment of Parkinson’s disease [28]. Finally, a safe verification process is needed to confirm the functional operation of the motion-adaptive algorithm in each patient.

### 2.2 Sensing and Classification

Inputs of the adaptive algorithms include time- and inertial-based signals. For time-based, circadian-synchronized control, the system scheduler uses an embedded real-time clock to send control signals to the microprocessor when a transition might be required. Using the clinician programmer, the scheduler can be configured to change the stimulation pattern according to the patient’s daily routine and circadian symptoms. The inertial sensing is provided by an embedded ADXL346, an ultra-low power microelectromechanical system (MEMS) three-axis accelerometer manufactured by Analog Devices [34]. The classifier leverages the digital motion processor (DMP) embedded in the ADXL346. The DMP is configured through the clinician interface through a read/write register field. While this interface requires referring to the register table in the manufacturer-provided documentation [34] to fully utilize, it does provide full accessibility to the DMP, which was deemed desirable for research teams exploring custom algorithms. The sensing axis, combination of axes, thresholds, AC/DC coupling, and timing constraints for rules/threshold-based classification provided by the DMP are all accessible in the register field.

To lower the programming burden, a set of reference register tables is provided to facilitate DMP configuration using representative use cases for algorithms based on absolute posture, general activity/inactivity, and transient shocks. Reference settings can be easily established and validated for various use cases, without requiring patient interaction, by using a helmet fitted with a digital twin of the part of the system that monitors and process the 3-axis acceleration and detects inertial events, as described in the supplementary material.

### 2.3 Control Policy: Integrating Circadian and Motion-Adaptive Algorithms

The control policy is implemented by allowing the circadian scheduler or the DMP to change the stimulation program by raising event signals (interrupts) in the embedded microcontroller. In the current realization two event signal lines are made available to the DMP. The two signals can be dynamically mapped to any two of single tap, double tap, activity, inactivity, free fall, and posture events in the register configuration (Fig. 3a). DMP events are used to select between two pre-configured stimulation programs, with their association configured on the clinician’s tablet programmer. In addition to the DMP-driven stimulation programs, the clinician also sets the default fallback program (per risk management) for the device. The final control policy constraint is to ensure that the stimulation amplitude ramps during program transitions are acceptable to the patient; the ramp rate represents a user-controlled trade-off between response time and side-effects, such as paresthesia. Given [35] the multiple control sources, we also needed to define the priority of events received by the stimulation controller (Fig. 3b). Based on our analysis of use cases, we chose to use the latest event arising from either the motion classifier, or manual intervention to determine the stimulation state. In addition to avoiding any confusion about prioritization, this approach allows for an intuitive hierarchy of expected changes: fixed patterns of stimulation from the scheduler are overwritten by more frequent motion-based therapy fine-tuning, while manual intervention will always override these automated adjustments, including disabling them completely. Fig. S4 shows an example of interaction between the scheduler and the motion classifier.

**Fig. 3.**
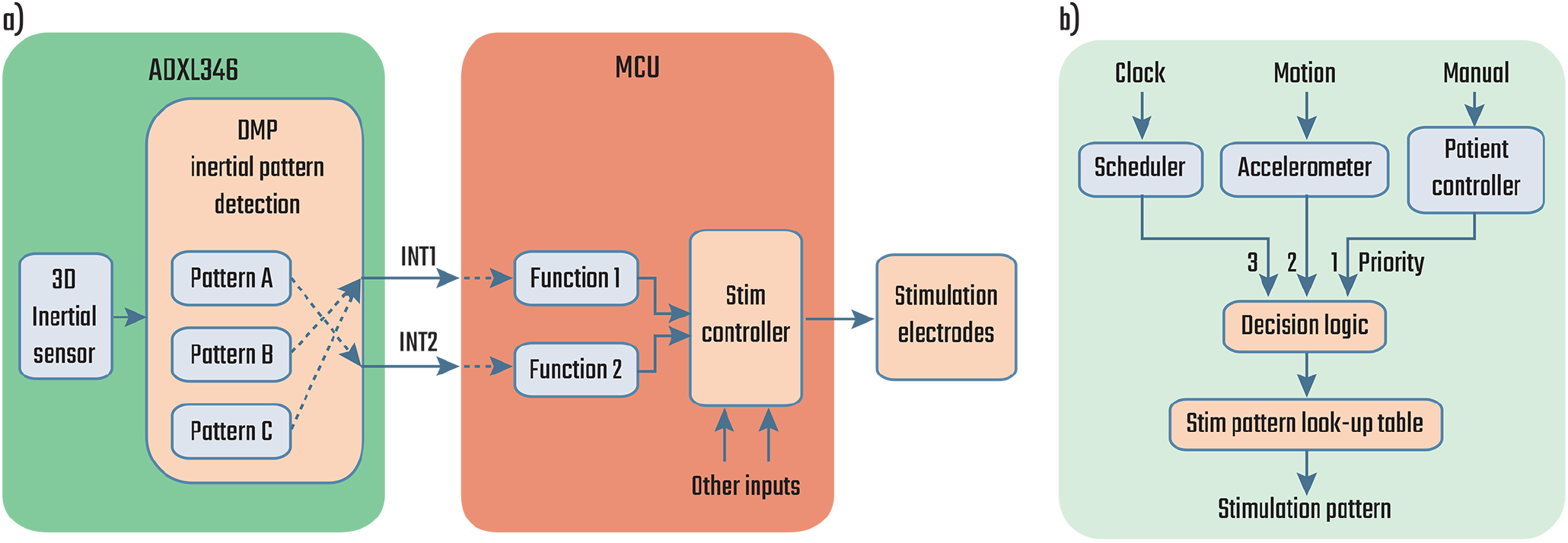
Block diagrams showing **a)** the pathway for the event signals (interrupts) generated by the DMP and **b)** the dynamic stimulation controller with its three types of inputs in increasing order of priority: fixed or scheduled, inertial detection, and manual control.

### 2.4 Risk Mitigations for Adaptive Systems: Actuation Limits, Fallback Modes, and Neural Dynamics

Using the physiologic control framework (Fig. 1), we identified potential hazards and specified systematic mitigations. We followed the ISO 14971 risk management process to identify and address potential harms to the patients. Particular emphasis was placed on the automated algorithms, and the IEC 60601-1-10 standard was used as guidance for the design of the control system [1]. With an automated system, the stimulation parameter space needs to be constrained to known-safe levels as the algorithm commands state changes. This “actuation limit” can be achieved by limiting the algorithm’s access to specific pre-configured programs (patterns of stimulation) [27, 36]. The clinician-researcher then effectively defines a boundary on parameters, with assurance that the algorithm never exceeds these limits. It is vitally important to provide visibility of the device state to the users, both the subject and the clinician. This observability was implemented on the patient controller with specific state alerts, including both the state of the algorithm (enabled/disabled) and the active stimulation program. All available states were also verified in software testing. Aligned with this specification, the patient controller also provides a mechanism to enable and disable the adaptive algorithm with a button press. Supporting the deactivation feature and stimulation limits, a preselected open-loop “fallback” program is also defined, which the stimulator defaults to upon manual termination of the algorithm [36]. Temporal safeguards on the algorithmic adjustments were also added including ramped transitions in intensity between stimulation programs to avoid subject discomfort such as paresthesia [37], and timing interlocks to avoid inadvertent rapid transitions at classification boundaries. As an abundance of caution, we specified that the adaptive motion algorithm should be disabled during recharge to prevent changes in the stimulation program and ensure a known stimulation state is always maintained throughout the process.

### 2.5 Acute Verification Methods: State Monitoring and Alerts

Once the motion-adaptive algorithm is configured, the verification of the automated system is supported through wireless telemetry to the patient programmer. When telemetry is enabled, the programmer interface is updated to display the implant’s embedded classification state. As the patient changes their motion state, the clinician-researcher can verify that the expected stimulation program is activated by monitoring updates telemetered to the patient controller.

## 3 System Verification

System verification ensures that the DyNeuMo Mk-1’s motion adaptive algorithms have provided the desired automated stimulation adjustments, while not compromising the existing functionality of the Picostim. A significant amount of the system hardware and software leverages the Picostim predicate, which allowed us to use existing verification testing protocols and reports for functional areas such as stimulation, telemetry, and biocompatibility.

The DyNeuMo Mk-1 verification efforts focused on the incremental additions of the accelerometer, adaptive algorithms, integration of circadian- and inertial-based inputs, and risk mitigation methods. Verification protocols demonstrated that the ADXL346 registers could be programmed appropriately for detection of specific inertial and activity states, and that stimulation was then adjusted accordingly. For example, Fig. 4 shows a representative state change that occurs when a subject becomes active (at time *t*_1_) or inactive (at time *t*_2_). Inertial transition points, timing interlocks, stimulation program mapping, and ramp rates were verified to operate as expected. Note that temporal responsiveness is fully programmable, as an example when a subject is laying down, the DMP could wait for several minutes to avoid symptoms while transitioning to sleep; however while standing up it could respond immediately to prevent falls. Other verification examples included activity vs inactivity (e.g. for essential tremor control or gait detection) by testing the AC-coupled accelerometer mode for classification. Finally, we verified tap/shock detection, which could be useful for transient events such as those related to urinary incontinence, or as a mechanical patient input that eliminates their need for interaction using the handheld controller. The stimulator can respond in under 15 ms to a transient event (Fig. S3), which falls within the reported acceptable latency for responding to mixed incontinence stress events [31]. The stimulation will stay active until the timing threshold for inactivity is met; in this demonstration, one second. Also verified was double tap detection, which can help improve classification specificity by reducing the likelihood of false positives.

**Fig. 4.**
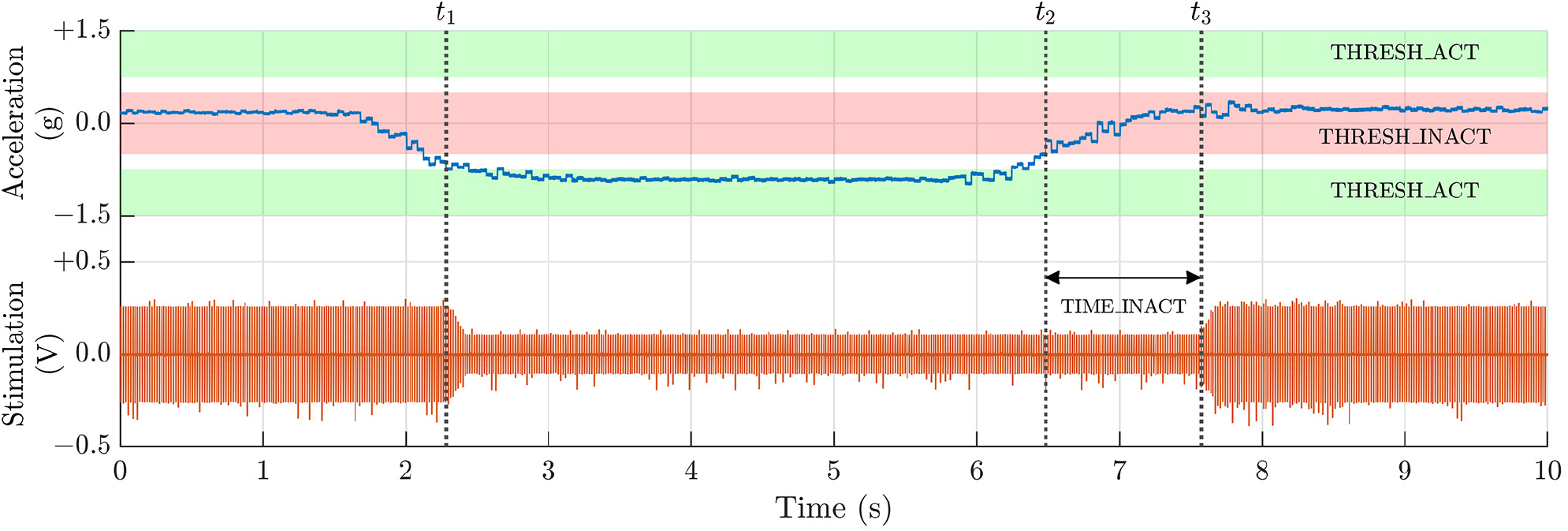
Data from the posture algorithm verification showing a stimulation transition including transition ramps and adjustable response timing. Note that activity response is set to be immediate (*t*_1_), while registering inactivity requires acceleration staying within the inactivity band for an adjustable interval *t*_2_ *-t*_3_ (nominally one second for demonstration purposes). Note the ramped transitions of stimulation intensity, a safety measure to avoid subject discomfort [37].

In addition to the functional performance of our adaptive algorithms, we also verified other system requirements such as power consumption, patient interface controls, and the human factors for algorithm programming. The power consumption of the MEMS sensor, including classification, is approximately 40 µW, or 10% of the nominal therapy for a Parkinson’s or essential tremor patient. Note that this estimate does not include any potential energy savings by turning down stimulation at night or during periods of low activity. Other key performance results are summarized in Table 2.

**Table 2.** Technical specifications for the DyNeuMo-Mk1 Investigational Research Syste

|  |  |
| --- | --- |
| **Sensor Characteristics** |  |
| Inertial sensing | 3-axis accelerometer, sensitive to 4 mg activity variations; dynamic range programmable $\pm 2$ g to $\pm 16$ g; typical sampling rate is 50 Hz |
| **Stimulation Characteristics** |  |
| Channel Access | 8 independent electrodes, typ. arrangement is 2 leads × 4 electrodes |
| Stimulation overview | 2 independent channels, current-controlled, charge-balanced monophasic / symmetric biphasic with programmable interphasic delay |
| Multiplexing | Full matrix configuration across electrodes (inc. the case reference) |
| Stimulation magnitude | 0–15 mA (0.05 mA increments) and 0–450 μs pulse width, fractionalized distribution available for guarded cathodes, etc; programmable and independent ramp rates |
| Recharge characteristics | Programmable passive and active recharge, with variable recharge ratio |
| Stimulation frequency | 1–500 Hz frequency stimulation; can go sub-Hz with cycling enabled; independent frequencies available across stimulation channels |
| Stimulation cycling | Adjustable cycle timing for enabling burst stimulation |
| Stimulation programs | Up to 8 independent programs can be configured in the IPG and accessed by algorithms or patient controller |
| **Algorithm Characteristics** |  |
| Motion Classification | Absolute orientation, activity vs non-activity (parameterized), shocks and free-fall detection |
| Stimulation Control policy | Detected classification states mapped to pre-configured stimulation programs with pre-specified transition ramp rate. Two independent stimulation programmes tied to motion states, and a default fall-back. |
| Risk mitigations | Algorithm implementation aligns to 60601-1-10 specifications for physiologic control loops (e.g. limits, alerts, data logs, fallback modes) |
| **Other System Characteristics** |  |
| Battery capacity, recharge cycle | < 2 h recharge with a target recharge interval of 4 days for typical Parkinson's-like settings |
| Mechanical Characteristics | Cranial-mount, 7.4 cc titanium package, with 2 leads for 4 contacts/lead |
| Telemetry/External Sensor and Stimulation Synchronisation | MICS-band radio, > 1 m distance, with hand-held module |
| MRI compatibility (in process) | MRI conditional imaging for 1.5 T and 3 T imagers |
| Electrodes | Modular design with ability to customize lead length and electrode spacing; currently supports brain stimulation and peripheral extradural electrodes and cuffs |
| Clinician programmer | Standard consumer tablet running Windows 10 |

## 4 System Validation in Movement Disorders: Cervical Dystonia

We consider system validation to be addressed through research protocols targeting specific disease states. To facilitate these experiments, the DyNeuMo Mk-1 is being released as an investigational research tool for the clinical neuroscience community including the design history files required to support investigational device approvals. In line with our user requirements, we aim to support existing therapies that might benefit from motion-adaptive stimulation; if the algorithm is not successful, it can be disabled and the patient still benefits at a minimum from the predicate therapy. We describe our pilot validation case here.

### 4.1 Case Description

The patient is 63-year-old woman with a five year history of cervical dystonia who underwent bilateral DBS implantation due to severe disability and poor quality of life, despite repeated botulinum toxin treatment. Her surgical procedure involved double targeting of the subthalamic nucleus (STN) and the ventralis oralis posterior nucleus (VoP) using a linear octapolar electrode (Boston Scientific) connected to a rechargeable implanted pulse generator. Dystonic head postures and pain improved considerably in the first two months post implantation. A progressive reduction of benefit then followed with persistent head torsion while walking. This resulted in renewed disability as the patient once again became unable to walk outside or perform household chores independently. Attempts to reprogram her DBS therapy settings involved the activation of different electrode contacts. Separate targeting of the left STN or VoP was effective in improving dystonia when walking or sitting down respectively. However, their combined activation failed to improve dystonic postures or pain and caused speech disturbances. Stimulating the right STN or VoP improved dystonic pain and head posture less effectively than the left side targets. Finally, the patient experienced rapid habituation to stimulation, with each setting alteration relieving symptoms for less than 24 hours.

### 4.2 Methods and Results

We hypothesized the patient required stimulation in two different left hemispheric targets to achieve full control of her symptoms when sitting down or walking. Accordingly, in July 2020 she was offered to be evaluated with a DyNeuMo Mk-1 device which could switch between stimulation programs based on motion state, potentially effective in controlling dystonia both when sitting down or walking. It was further hypothesized that regular changes to stimulation program from everyday activity would help in preventing habituation to therapy. Our aim was to use the results of this intervention to optimize the patient’s long-term care with an existing CE–marked system. The evaluation received humanitarian exemption authorization from the Medicines and Healthcare Products Regulatory Agency (MHRA), and the patient provided signed written consent to be trialled with an externalized DyNeuMo Mk-1.

The DyNeuMo Mk-1 accessed the patient’s existing implanted leads and extensions through a disposable extension adaptor, as presented in Fig. 5. The device was placed over the cranium to represent the approximate intended location of a typical DyNeuMo system. Stimulation parameters were optimized for different motion states, and the classifiers were set to detect these states. On the left side, the device was set to switch between a “sitting down/standing up” program (employing a contact in VoP, 5-) and a “walking” program (employing a contact in STN, 3-). On the right side, the best contact for both sitting and walking was localized in VoP (4-). All stimulation electrodes were returned to the case through a conducting electrode pad attached to the skin of the shoulder. The control policy algorithm mapped the appropriate stimulation program to the classified motion state, with a ramp rate programmed (nominally 0.2 mA/s) to avoid side-effects.

**Fig. 5.**
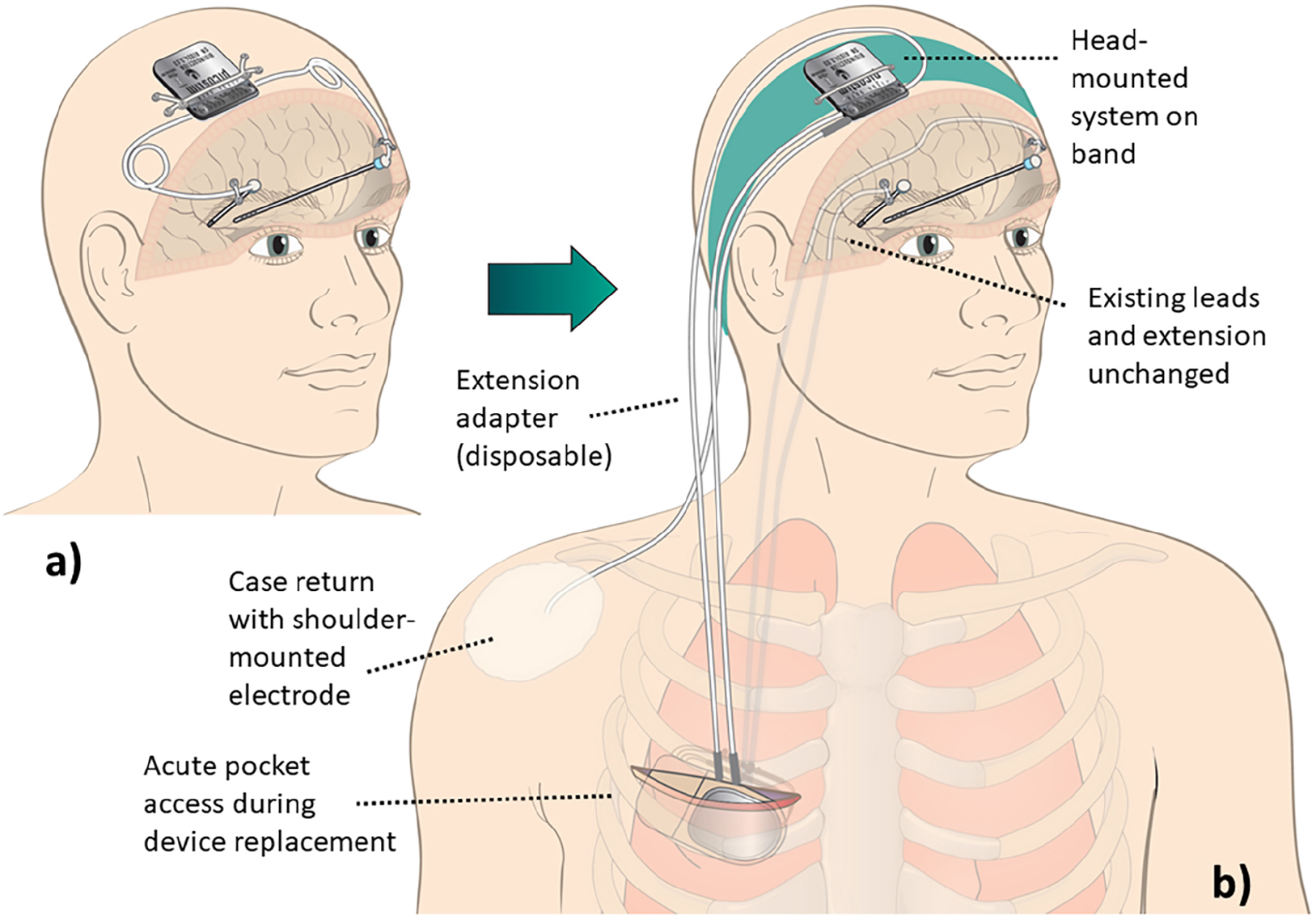
**a)** Intended use case for the DyNeuMo Mk-1 for deep brain stimulation. **b)** Validation experiment set-up. The patient’s implanted DBS lead extenders were exposed, at the surgical pocket formed during device changeout. The implanted lead extenders were connected to the externalized DyNeuMo Mk-1 through custom-made disposable DBS lead adapters outside the body. For the duration of acute testing, the commercial replacement IPG was left unconnected. The DyNeuMo was placed over the cranium using a wearable headband to be representative of its intended point of implantation. For the return electrode, a conducting electrode patch was placed on the shoulder and connected to the case. This system configuration allowed for subacute testing during device change-out.

The efficacy of the adaptive mode was then evaluated. When tested without stimulation, the subject scored 16 out of 25 on the modified Tsui scale rating for cervical dystonia severity. With stimulation and the adaptive algorithm enabled, the patient showed marked and immediate improvement (within 30 seconds) of symptom severity both when sitting down and while walking, scoring 2 out of 25 the modified Tsui scale. The benefits persisted at 36-hours assessment, with no evidence of habituation.

No significant adverse events were observed. The study was carried out over six days; four days were spent on configuring and testing the motion adaptive mode and adjusting the stimulation parameters, followed by two days of testing with the best parameter set. The stimulation parameters were optimized for two motion states using the DyNeuMo Mk-1 to both control the stimulation program and detect the change in motion state. As the patient’s replacement device lacked motion adaptive capabilities, it was configured with a set of parameters that represents a compromise between the best set of parameters for each motion state found with the DyNeuMo, allowing for informed, evidence-based protocol selection.

## 5 Limitations of the DyNeuMo Mk-1 adaptive algorithm

The DyNeuMo Mk-1 does have significant limitations worth noting; these are both technical and physiological. Perhaps most importantly, the current embodiment limits the measurement of motion to the device implantation site. In the case of a cranially-mounted system such as the predicate Picostim for deep brain stimulation, the specific measurement of hand tremor is therefore not supported; a more general correlation with general motion is required, which limits the specificity of the adaptive algorithm. An additional specificity error arises from the measurement limitations of a three-axis accelerometer. Specifically, the DMP can be confounded when estimating posture by the superposition of linear acceleration with the gravitational field. This concern can be addressed somewhat by adjustment of the time and level constraints before generating an event signal, but this mitigation is a trade-off with transition latency. While a gyroscope might help improve specificity, it also requires significantly more power than permissible within the power budget of most stimulation therapies due to the principles of MEMS– based Coriolis sensing; if the application allowed for it, duty-cycling might help somewhat resolve this issue. Finally, our setup is currently constrained to only two motion-based stimulation states. If this is found to be severely limiting, we could perform more advanced event masking and explore adaptive DMP register adjustments in the future. As an example of physiological limitations, the time dynamics between stimulation and physiological response need to align with the adaptive algorithm capabilities. For example, if stimulation requires extended time to exert therapeutic effect, then the utility of adaptive stimulation titration might be limited. At this time, we believe that several clinically-meaningful adaptive algorithms can be implemented with the first generation research tool, and we can refine future designs based on relevant clinical experience.

## 6 Conclusion

There is growing interest in adaptive medical devices to improve therapies by automatically adjusting stimulation based on clinically-relevant physiological features. We have developed a fully-implantable medical device that integrates circadian and inertial signals as two sources physiological inputs. The advantages of our approach are that it is 1) relatively easy to configure the classifier for clinically-relevant states, 2) relatively inexpensive to manufacture, and 3) highly reliable as a method. Integrating two input sources, with significantly different temporal dynamics, created unique design issues for prioritization and risk mitigation. However, the complete algorithm ultimately reflects the dynamics found in many physiological control processes balancing circadian (feedforward) and homeostatic (feedback) constraints.

The integration of circadian and inertial sensing could be a practical solution for several unmet needs, and was validated in our subacute case in cervical dystonia. In other validation cases, we are exploring the treatment of orthostatic hypertension, gait imbalance, and sleep disturbances using deep brain stimulation of the pedunculopontine nucleus [38, 39]. Our choice of this protocol is motivated by the relationship between inertial signals, clinical state, and stimulation parameters that can be explored with motion-adaptive stimulation, while also being aware that stimulation of the reticular activating network can result in sleep-wake disturbances, which motivates the integration of circadian-based algorithm constraints. We recently reported on a case study in managing status epilepticus using the DyNeuMo system in a canine patient, supporting the utility of our combined circadian and activity-based therapy approach [40]. As for in-human research, the DyNeuMo Mk-1 has been approved for use in the commencing MINDS-MSA trial for the treatment of the motional symptoms of multiple system atrophy, while the predicate Picostim device is currently being evaluated in the SPARKS trial (NCT03837314) for use in Parkinson’s disease, and in the STAG-MSA trial (NCT03593512) for multiple system atrophy.

Pending promising outcomes from these trials, we hope to expand to other disease states where explicit mappings between inertial signals and desired stimulation exist, such as tremor, cervical dystonia, and urinary incontinence [22, 41]. Going forward, the second generation of DyNeuMo systems is in development to extend the control framework with bioelectrical sensing and classification for advanced therapy research [42].

## Author Contributions

Clinical contributors F.M., E.P., S.M., T.Z.A., and A.L.G. provided input on DBS therapy and neurosurgery, defining the user needs and target specifications. J.O., T.G., G.L. and I.G. worked on developing the Picostim platform, with T.N., F.M. and E.P. leading compliance and preparatory work for the human trials and emergency exemption case. T.G.C., and J.H. were involved in technology definition, review, and verification. T.D. supervised the DyNeuMo project and system-validation activities. M.B. worked on hardware verification. M.Z., Z.N., and D.S. configured and verified the motion classifier subsystem, with R.T. contributing analysis. M.Z., R.T., T.D. and F.M. prepared the manuscript, which all authors reviewed.

## Acknowledgements

The authors would like to thank Stuart M. Higgs for the design and assembly of an ADXL346 evaluation circuit, and Josephine Blyth for creating and testing the helmet-mount accelerometer reference setup.

