## supplementary material for "DyNeuMo Mk-1: Design and Pilot Validation of an Investigational Motion-Adaptive Neurostimulator with Integrated Chronotherapy"

### 1 Hardware validation

We developed a system for classifier training and verification. The training module included real-time data streaming at 10 Hz through the MICS-band radio, while logging the accelerometer data to a file that can be used to determine register values for a motion-specific classifier. An example of the real-time streaming is provided in Fig. S2 which is compared to an external reference accelerometer. As illustrated in Fig. S1, the training module includes a wearable sensor that acts as a twin of the implant. This system, which acts as a digital twin for the DyNeuMo Mk-1 motion adaptive classifier, allows for register settings and classifier outcomes to be established before committing to the implant. Full description of the system, including hardware schematics, microprocessor code, and graphical user interface have been archived and made available at doi: 10.5281/zenodo.5745253.

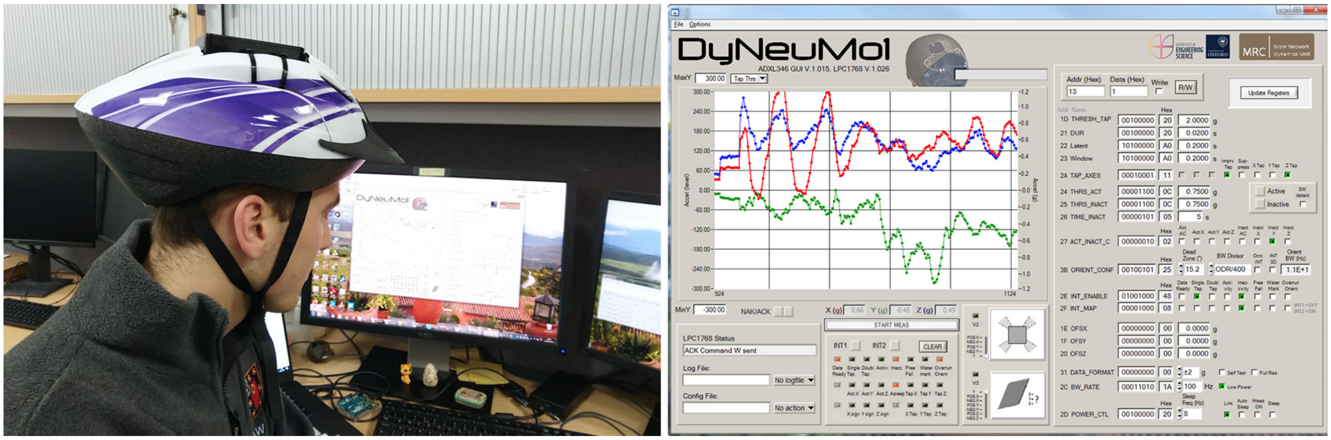


**Fig. S1.** Prototyping system for inertial measurements and classification. Left: the reference accelerometer can be mounted on different body locations for sensor measurements and classifier assessment; shown here on a bicycle helmet for assessing cranial placement. Right: user interface for displaying the signals, reading and writing registers, and assessing classifier outputs and interrupts.

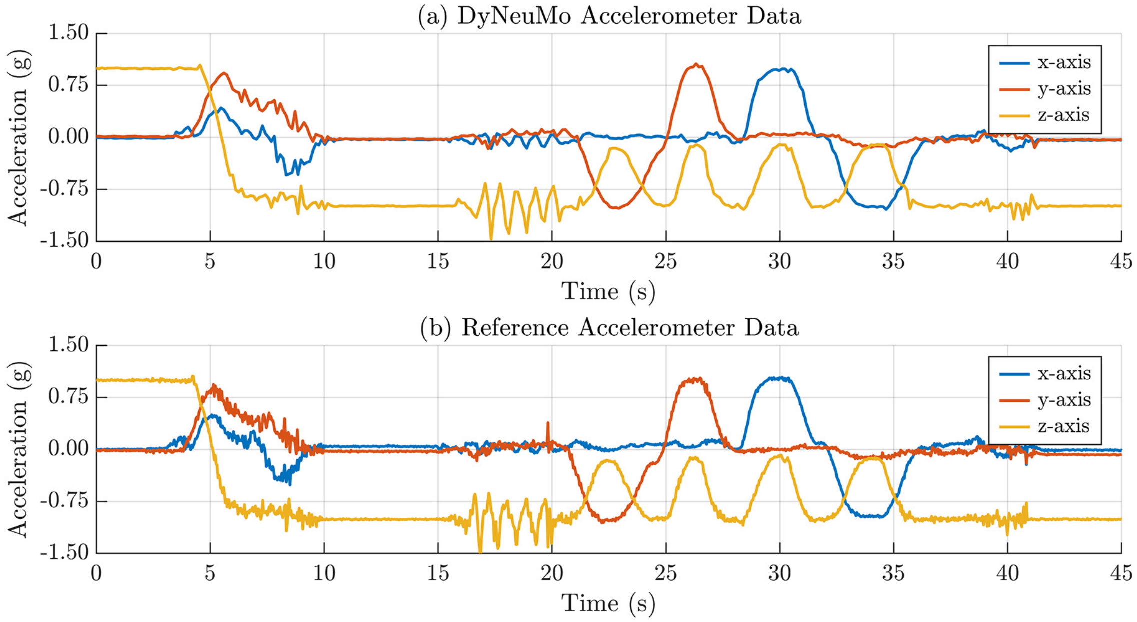


**Fig. S2.** Validation of the data stream used for gathering user specific training data for the classifier. **a)** Data streamed from the DyNeuMo Mk-1 implantable system through the MICS-band radio, shown with interpolation. **b)** Reference accelerometer data used for external calibration. Note that the embedded accelerometer closely follows the reference, albeit at a lower sampling rate.

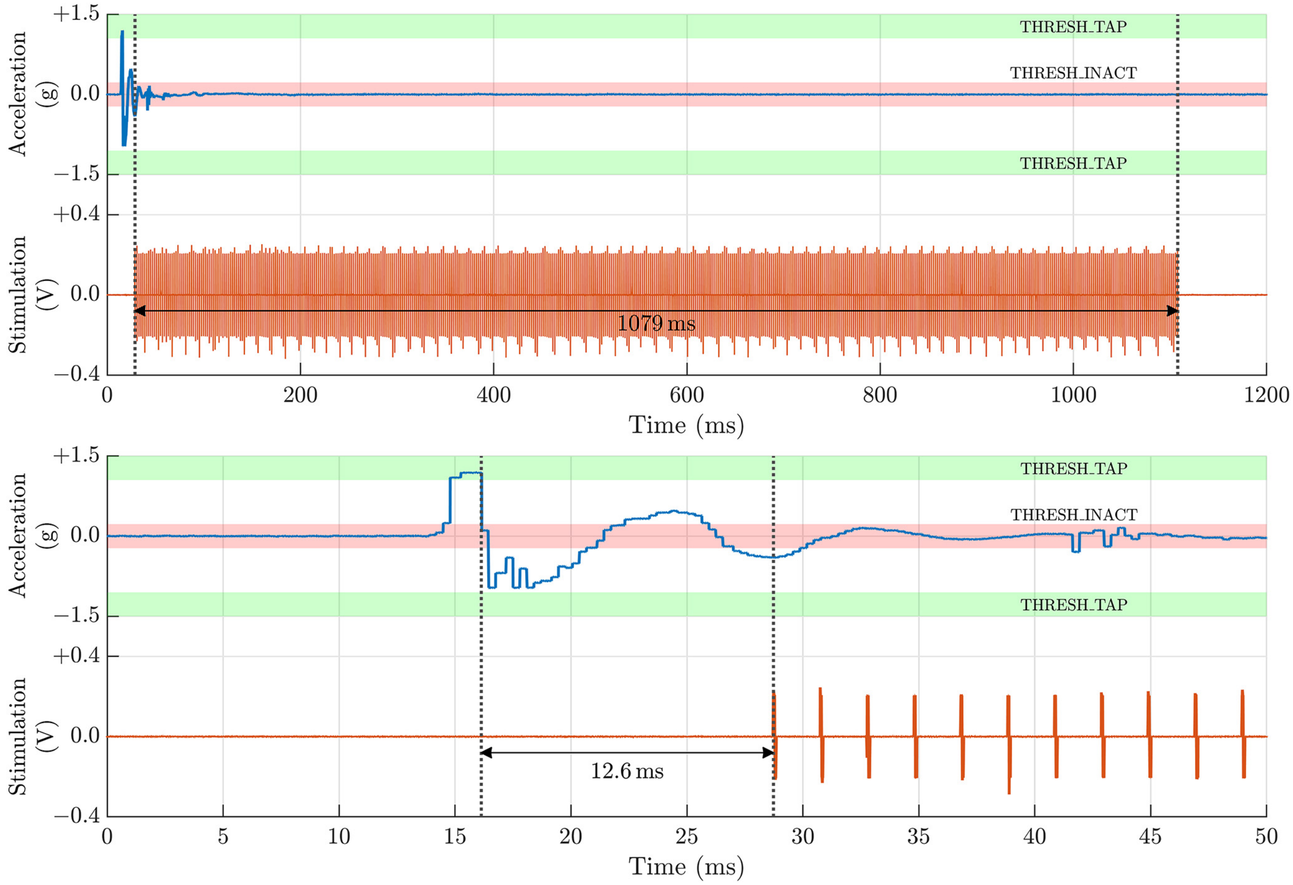


**Fig. S3.** Timing characterization for stimulation. Top panels: the initiation of stimulation due to a transient shock, that ceases after a programmed second of inactivity. Bottom panels: Resolving the response time to the shock, which shows stimulation initiating roughly 12.6 ms after detection.

### 2 DyNeuMo Mk-1 adaptive mode benchtop validation

This section includes audiovisual demonstration of the device adaptive mode for three use cases. The last video shows two of risk-mitigation measures, transition ramps and fallback mode.

**2.1 Walking / sitting or standing still.** Walking may be particularly challenging for Parkinson's disease patients, requiring an increase in DBS amplitude. On the other hand, patients receiving DBS therapy for other conditions may experience side effects that affect their balance. Therefore, it can be beneficial to be able to detect when the patient is walking and adapt the stimulation to either aid in the execution of the movement or reduce the side effects of the stimulation. In this demonstration, the DyNeuMo Mk-1 implantable pulse generator (IPG) is configured to detect two inertial states. One is walking, while the other is sitting or standing still. A pre-configured stimulation pattern has been assigned to each motion state.
`https://youtu.be/40_kKova5YU`

**2.2 Posture detection: horizontal / vertical position.** Patients with MSA usually suffer orthostatic hypotension. Adapting the DBS therapy to changes of posture could help to regulate the blood pressure of these patients according to their body posture. The video in the link shows the response of a DyNeuMo Mk-1 IPG that has been configured to detect a change of posture, either horizontal or vertical position. A pre-configured stimulation pattern has been assigned to each posture state.
`https://youtu.be/X1_FVKsif-I`

**2.3 Single tap / double tap.** The patient or the carer can change the pattern and amplitude of the stimulation by using the DyNeuMo Mk-1 patient controller (Picon). In some situations, it could be more convenient and discreet to tap on the device with the fingers to change the stimulation pattern. This could be particularly useful in an emergency situation, for instance, a seizure in an epileptic patient. In this video, the DyNeuMo Mk-1 IPG is configured to detect two inertial states: single or double tap. A pre-configured stimulation pattern has been assigned to each state.
`https://youtu.be/vxG7_czmdIk`

**2.4 Transition ramps and fallback mode.** The DyNeuMo Mk-1 IPG is configured to have a smooth transition when the stimulation amplitude changes. When the adaptive mode is turned off, the device switches to a pre-configured, safe stimulation pattern as a fallback mode.
`https://youtu.be/26_L0j7wsVs`

### 3 Interaction between the scheduler and the adaptive mode

Fig. S4 illustrates how the DyNeuMo Mk-1 handles the signals from the circadian scheduler and the adaptive mode. The schedule, configured by the clinician using the programmer interface (top), covers the 24 hours of the day with a 30-minute resolution. In this figure, pre-configured daytime (red) and night-time (blue) stimulation patterns are fed into the daily schedule of the patient. The use case presented here is of a patient with generalised epilepsy. The patient seizure diary shows that the majority of the seizures occur during sleep. Therefore, the DMP in the device is configured to detect inactivity after acceleration in all three axes have fallen under a threshold for more than 1 minute and 15 seconds. When inactivity is detected, the stimulation changes to a pattern with a higher amplitude (amber). The stimulation pattern goes back to the next scheduled pattern once the patient wakes up and the acceleration in any of the axes goes above the inactivity threshold. Additionally, the patient's carer can activate the emergency or boost stimulation pattern (green) when a seizure occurs by a single tap on the device using the fingers on the patient's scalp. In most cases, the boost pattern will halt the seizure and the stimulation goes back to the next scheduled pattern. Otherwise, the carer will use the manual controller to disable the scheduler and maintain the boost pattern for as long as it is required.

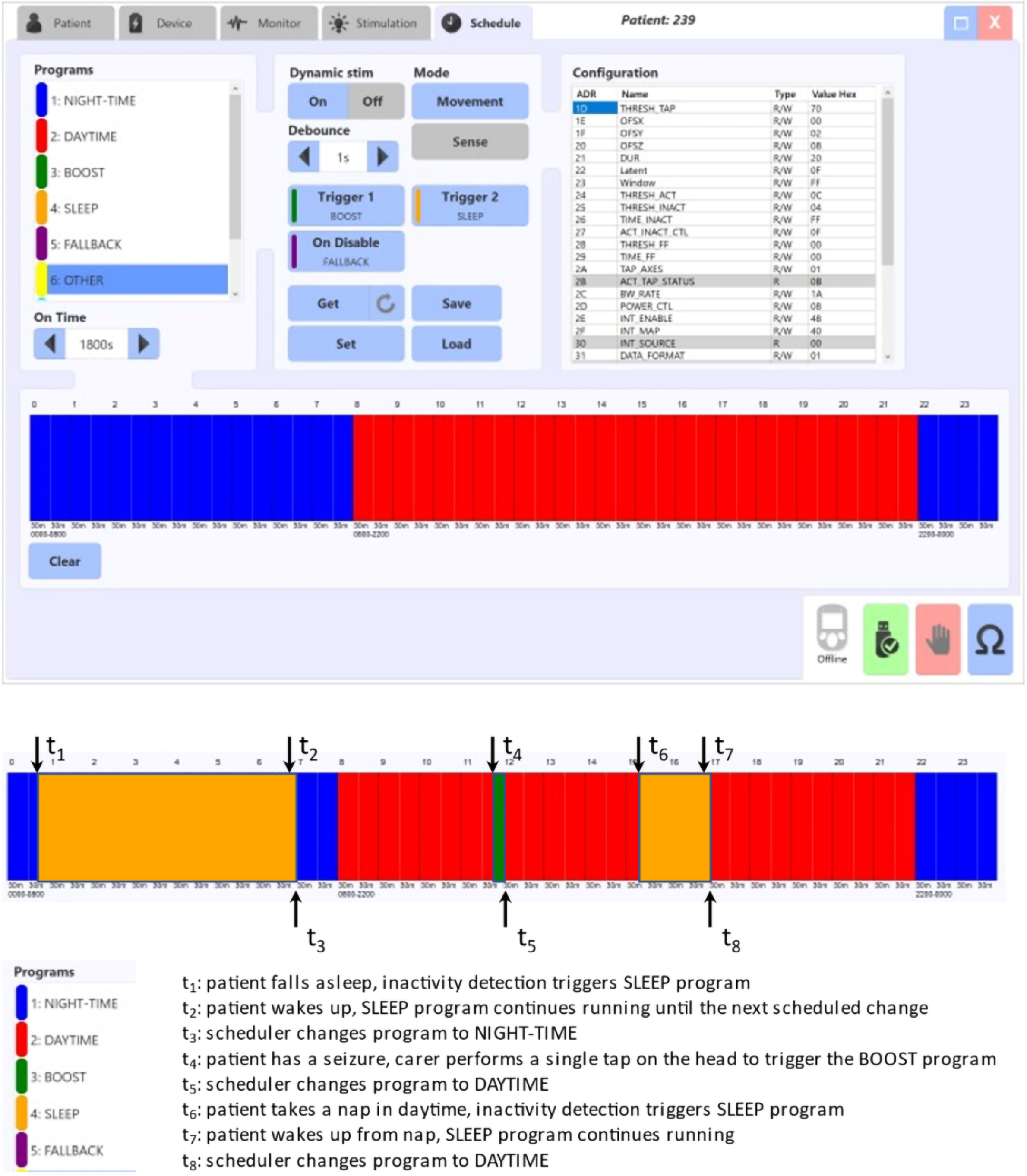


**Fig. S4.** Interaction between the circadian scheduler and the adaptive mode.
